# Cholinergic modulation supports dynamic switching of resting state networks through selective DMN suppression

**DOI:** 10.1101/2022.11.01.514686

**Authors:** Pavel Sanda, Jaroslav Hlinka, Monica van den Berg, Antonin Skoch, Maxim Bazhenov, Georgios A. Keliris, Giri P. Krishnan

## Abstract

Brain activity during the resting state is widely used to examine brain organization, cognition and alterations in disease states. While it is known that neuromodulation and the state of alertness impact resting-state activity, neural mechanisms behind such modulation of resting-state activity are unknown. In this work, we used a computational model to demonstrate that change in excitability and recurrent connections, due to cholinergic modulation, impacts resting-state activity. The results of such modulation in the model match closely with experimental work on direct cholinergic modulation of Default Mode Network (DMN) in rodents. We further extended our study to the human connectome derived from diffusion-weighted MRI. In human resting-state simulations, an increase in cholinergic input resulted in a brain-wide reduction of functional connectivity. Furthermore, selective cholinergic modulation of DMN closely captured experimentally observed transitions between the baseline resting state and states with suppressed DMN fluctuations associated with attention to external tasks. Our study thus provides insight into potential neural mechanisms for the effects of cholinergic neuromodulation on resting-state activity and its dynamics.

## 1 Introduction

Resting-state fluctuations have been established as one of the fundamental properties of spontaneous brain dynamics observed across different species in neuroimaging studies [Biswal et al., 2010, Hutchison et al., 2011, Chuang and Nasrallah, 2017]. However the neural mechanism underlying the origin of the fluctuations remains poorly understood. High amplitude fluctuations are the main contributor to emergent patterns of functional connectivity patterns which are used to define the underlying architecture of functional networks [Allan et al., 2015, Esfahlani et al., 2020]. Analysis of these networks received a great deal of attention [Van Den Heuvel and Pol, 2010] and connectivity changes in disease states are considered to reflect underlying pathologies [Fox, 2018].

One of the most intriguing observations is the increased spontaneous activity in specific brain areas during rest periods. Early observations identified the Default mode network (DMN) as a sub-network that has higher activation and functional connectivity during rest periods compared to task periods (Fox et al. [2005], Raichle and Snyder [2007]). The DMN is now regarded as a fundamental functional network activated during internal processing modes and deactivated when attention shifts to external tasks [Buckner and DiNicola, 2019].

What governs the transitions between DMN-dominated rest states and DMN-suppressed attentive states is widely speculated but not yet clear [Sridharan et al., 2008, Ghosh et al., 2008, Chen et al., 2013, Hansen et al., 2015]. One hypothesis states that chemical neuromodulation plays a critical role in dynamic transitions between functional networks [Li et al., 2015, van den Brink et al., 2019, Shine, 2019]. Specifically, cholinergic activity has been suggested to influence the balance between rest and task-related brain activity [Hahn et al., 2007, Tanabe et al., 2011, Sutherland et al., 2015]. The basal forebrain (BF) is a major source of acetylcholine (ACh) in the neocortex with broad yet specific projections [Zaborszky et al., 2015b, Záborszky et al., 2018, Markello et al., 2018]. Activity in BF closely matches the activity of the default mode-like network (DMLN, animal homologue of DMN, Nair et al. [2018]) in animal models, and changes in co-activation of BF and DMLN was reported in early-stage rodent models of Alzheimer’s Disease [van den Berg et al., 2022]. In fact, BF itself was suggested to be an inherent part of DMN [Alves et al., 2019, Lozano-Montes et al., 2020]. In a recent work from our group, we measured changes in resting-state fMRI following exclusive activation of BF cholinergic neurons in transgenic ChAT-cre rats using chemogenetics. In particular, the injection of a synthetic drug (CNO) designed to activate DREADD receptors expressed on BF cholinergic neurons resulted in decreased spectral amplitude and functional connectivity in DMLN, a hallmark of DMN suppression [Peeters et al., 2020]. While these studies provide evidence for cholinergic modulation of resting-state activity, the underlying neural mechanism has not yet been identified.

In this study, we identify a possible mechanistic explanation for the cholinergic modulation of resting-state activity. We used a large-scale biophysical *in-silico* network model of rat and human connectome. The advantage of cellular-level biophysical modeling is that we could model the influence of ACh neuromodulation as changes to ionic and synaptic currents. Cholinergic modulation is not fully understood and vary by cell type and region [Colangelo et al., 2019]. Here, we identified that excitability and recurrent connection as potential critical components of cholinergic modulation that impacts resting-state activity. We used experimental results of direct cholinergic neuron manipulation to constrain our model of the DMLN network. We first demonstrate that the computational model can capture the same changes in DMLN as observed during cholinergic modulation in rodents. We then extend these findings using a computational model of human connectivity derived from diffusion-weighted MRI (DW-MRI). Finally, we demonstrate that a selective increase in cholinergic activation only in DMN results in the suppression of DMN activity and functional connectivity without modifying sensory networks.

## 2 Results

We used a biophysical model of infra-slow resting-state fluctuations based on our previous work [Krishnan et al., 2018]. This model includes a network consisting of conductance-based excitatory and inhibitory neurons with realistic synaptic AMPA, NMDA, and GABA synaptic connections. In addition, dynamic variables corresponding to intra and extracellular ion concentrations, including K^+^, Na^+^, Cl^-^, and Ca^2+^ions, were included. In this model, the slow variation of ion concentration in time, specifically extracellular K^+^ion concentration, allows for slow change in the neuron’s excitability and firing rate, leading to fluctuations in the 0.02 Hz range similar to slow resting-state activity (Fig. 1D-F). In this current work, we extend our previous model to incorporate cholinergic modulation through direct manipulation of intrinsic and synaptic currents.

**Figure 1:**
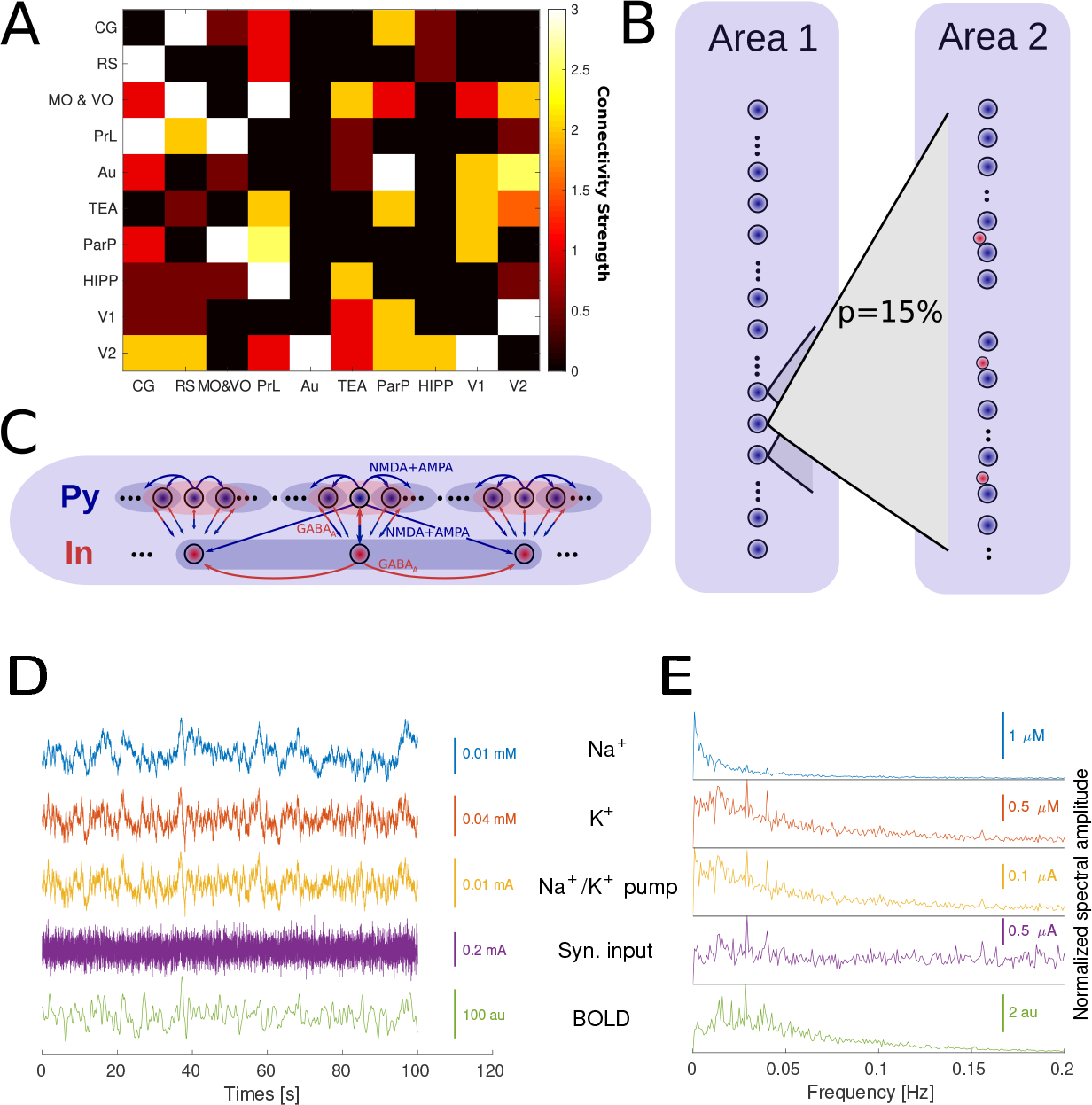
A. Structural connectome of a rat’s DMLN (imported from neuroVIISAS project). B. Model connectivity between two distinct DMN areas. Each source neuron has a p=15% probability of connection to each neuron in the target area. The connectivity strength of AMPA connection between excitatory neurons was derived from the structural connectome in A. C: Model connectivity of excitatory and inhibitory neurons within a single module (area). D: Example traces of Na^+^and K^+^extracellular levels, the activity of Na^+^/K^+^pump, average dendritic excitatory synaptic input (average across 50 excitatory neurons in a single area and 100 ms time window) and resulting BOLD trace (in arbitrary units). E. Spectral amplitude of the same variables as in E (single trial, average across all areas).

Past experimental work has shown that acetylcholine modulates several ionic currents and synaptic currents through nicotinic and muscarinic receptors. Based on these findings, we implemented a detailed model of cholinergic modulation as a reduction of somatic and dendritic potassium leak currents 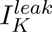, somatic delayed-rectifier potassium current *I_Kv_*, slowly activating potassium M-channel current *I_Km_*, high-threshold Ca^2+^ current *I_HV A_*, Ca^2+^–sensitive K+ current *I_KCa_*in excitatory neurons based on past experimental work [McCormick and Prince, 1986, McCormick, 1992, McCormick et al., 1993, Galvin et al., 2020]. In inhibitory neurons, the somatic and dendritic 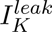 current and somatic *I_Kv_* current were reduced. Further, the influence of ACh on synaptic transmission was implemented as a decrease in excitatory AMPA connections and increase in NMDA connectivity based on experimental work [Gil et al., 1999, Hsieh et al., 2000, Vijayraghavan et al., 2018]. By examining the impact of change in each of the ionic and synaptic currents on resting-state activity, we identified that K^+^leak and AMPA current had the largest impact on resting-state activity. This led us to use a simplified model with changes to only K^+^leak and AMPA currents for large-scale network simulations (see the next section).

For the network simulations, the connectivity between different brain regions was identified through structural imaging methods: DW-MRI (humans) and axonal tracing (rodents). On the global scale, the brain regions were connected with diffuse long-range connections (Fig. 1B). The strength of the connections was proportional to the weight between regions of the global structural connectome (Fig. 1A). On a finer scale, each modeled brain region was represented by a locally connected group of 50 excitatory neurons and 10 inhibitory interneurons (Fig. 1C). In the case of rodent brain simulations, the connectome representing the structural connectivity of the rat’s DMLN was derived from the NeuroVIISAS database of axonal tracing studies [Schmitt and Eipert, 2012].

### 2.1 Cholinergic modulation of DMN resting state in rodents

To constrain and validate the effects of ACh modulations in our model, we used data from our previous chemogenetic experiments in transgenic ChAT-cre rats [Peeters et al., 2020]. This transgenic rat model allowed the selective targeting of cholinergic neurons in the BF (the primary source of ACh release to the neocortex) using designer receptors exclusively activated by designer drugs (DREADDs) [Alexander et al., 2009]. In these experiments, Blood Oxygenation Level Dependent (BOLD) activity was measured in the resting state before and after injection of CNO, a synthetic chemogenetic activator of DREADD receptors that were expressed exclusively in BF cholinergic neurons, resulting in widespread ACh release in the projections areas of these cells. To control for any off-target effects, CNO was also injected in animals which did not express DREADDs (sham animals). A second control, which we use as a baseline, was performed by injecting vehicle (saline) in the DREADD expressing animals with both control conditions resulting in no significant effects in the BOLD and functional connectivity. On the other hand, injections of CNO in the DREADD animals (ACh release) resulted in decreases in BOLD amplitude and spectral profile as well as reductions of functional connectivity and the fractional amplitude of low-frequency fluctuations (fALFF, Zou et al. [2008]) in DMLN (Fig. 2A).

**Figure 2:**
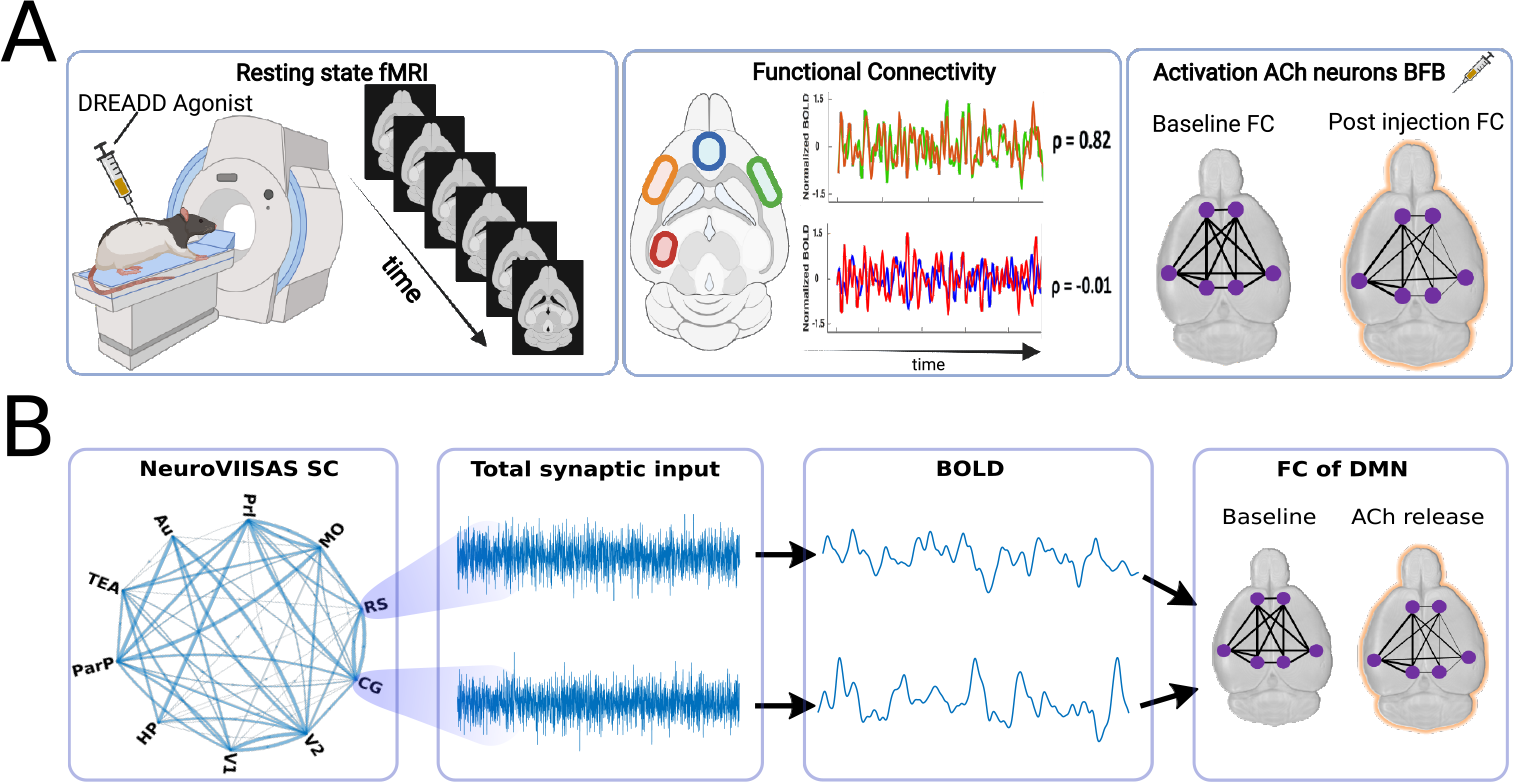
Experimental and modeling methods. A. Experimental framework. Chemogenetic tools (DREADDs) were used to selectively increase cholinergic activity in rat’s basal forebrain (BF). Resting-state fMRI scans were performed during the resting state/after injection of saline and after the injection of CNO, resulting in upregulated cholinergic release in BF. Functional connectivity and other signal features were collected to compare both conditions. B. Simulation framework. The structural connectome of DMLN is the backbone of simulation dynamics. The total synaptic input of all neurons within each area is measured and used for conversion to a BOLD signal. The correlation of BOLD between areas was used to define functional connectivity of DMLN in two conditions – resting saline baseline and increased cholinergic release.

We then used a computational framework that sufficiently mimicked the experiments. The connectivity of rat DMLN was used and synaptic activity from each anatomical area was used to derive BOLD signals, see Fig. 2B. We compared the simulation during baseline resting state and during high ACh levels that correspond to the experimental post-saline baseline and CNO conditions, respectively (Fig. 2B). We monitored several dynamic variables of the model – including firing rate, the activity of Na^+^/K^+^pump, extracellular K^+^levels, neuronal membrane voltage (spiking) and synaptic input. The total synaptic input was transformed into a BOLD signal representing the dynamics of each brain area (see Methods).

We first performed systematic variations of K^+^leak and AMPA conductance and identified that a reduction of 8% in K^+^leak currents and a 20% reduction in AMPA currents (local and long-range connections) matched the results observed in DREADD experiments in both detailed and reduced model (see SI Fig. 9). For the subsequent sections, we only present results from model involving only change to K-leak and AMPA conductance.

The amplitude of the BOLD signal was reduced during elevated ACh condition in the model and experimental recordings (compare the top and bottom panels of Fig. 3). The power spectrum in the low-frequency range (0-0.1 Hz) and fALFF measured across multiple trials were reduced with an increase in ACh. The experiment’s spectral profile tended to have a wider frequency range (0-0.15 Hz) compared to the model (0-0.1 Hz). This difference is partly due to lower variability in the peak of resting-state activity in the model compared to the experiment. The average Z-score, which measures functional connectivity between all the regions, was also reduced following an increase in ACh. Similar reductions were also observed in Na^+^/K^+^pump currents and fluctuations of extracellular K^+^concentrations. Overall, the change in excitability and recurrent connections due to ACh release in the DMLN model were sufficient to reproduce the changes in the BOLD signal of rat’s DMLN following selective cholinergic neuron activation in BF.

**Figure 3:**
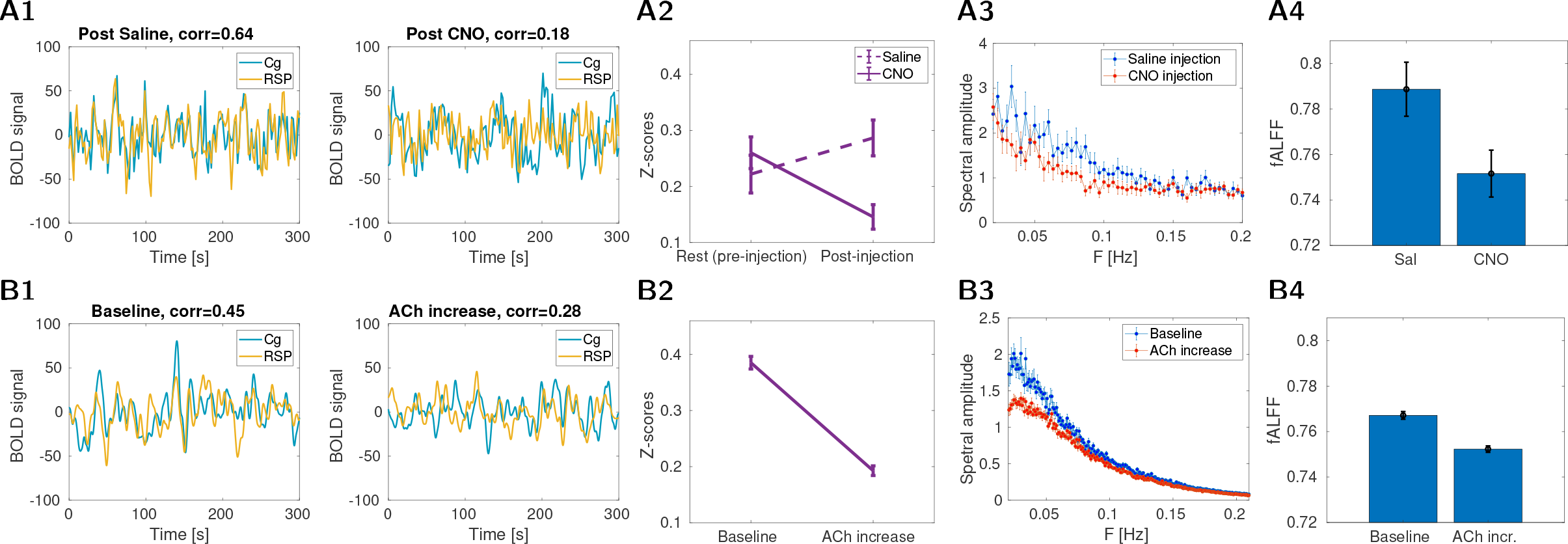
Top row: Properties of BOLD signal in rats. A1: Example traces of BOLD in the cingulate and retrosplenial cortex. Left: post-saline condition (“baseline”), right post-CNO condition (“ACh activation”). A2: Average functional connectivity comparing resting control and condition 15-20 min after DREADD saline/CNO injection (16 animals in each condition), Fisher z-transformed correlation *±*SEM. A3: Group-averaged power spectra of the seed-based maps of the cingulate in the right hemisphere (similar results can be obtained for the retrosplenial cortex). Blue curves are the power spectra after injection of saline; orange curves are the power spectra after injection of CNO. A4: fALFF values calculated from the seed-based FC maps of the right cingulate cortex (similar results can be obtained for retrosplenial cortex, see Peeters et al. [2020]). fALFF values were extracted from voxels of the right hemisphere after saline injection and CNO injection. The bar graphs present mean fALFF values *±*SEM. Bottom row: Properties of BOLD in simulations. B1: Example traces of BOLD signal in the cingulate and retrosplenial cortex. Left: Spontaneous activity of the model (“baseline”), right: activity in the condition of ACh release. B2: Average functional connectivity comparing baseline and ACh condition (16 trials in each condition), Fisher z-transformed correlation *±*SEM. B3: Average power-spectra from all DMLN areas (*±*SEM, 16 trials). The blue curve is the baseline condition, orange ACh condition. B4: fALFF values calculated from all DMLN areas. The bar graphs present mean fALFF values *±*SEM.

### 2.2 Resting-state activity in a large-scale model of the human brain

In order to examine the cholinergic changes in humans, we first establish a baseline computational model with realistic structural connectivity based on human DW-MRI. The structural connections for the whole brain model were derived from the dataset of 90 healthy subjects (see Methods for details), describing the structural connectivity (SC) among brain regions defined by the AAL atlas [Tzourio-Mazoyer et al., 2002]. While the extraction of structural connectivity matrix from DWI data generally faces a range of challenges [Schilling et al., 2019] and may depend substantially on the particular method used, the current structural connectivity data have been thoroughly quality controlled and validated against an independent dataset (see Škoch et al. [2022] for details). The resulting average structural connectivity matrix for a single hemisphere is shown in Fig. 4A1 (connectivity for both hemispheres and histogram of relative coupling strength between the brain areas is shown in SI Fig. 10). We observe typical properties of structural connectivity matrices extracted from DWI data. Namely, there is a skewed distribution of links with a small proportion of very strong links and a clustered structure with high density within functionally related brain areas. Further, the connectivity within the right and left hemispheres were very similar, giving rise to an almost symmetrical structure for the whole brain matrix. Because of this, together with the fact that inter-hemispheric connections are still imperfectly captured by current tractography methods, and higher computational demands, we used only a single hemisphere in our model and analysis (but see SI Fig. 10E for an overview including both hemispheres). We also measured 15 mins of fMRI measurements in resting conditions from the same human subjects we used to estimate the structural connectivity, allowing us to estimate functional connectivity. Fig. 4A2 shows the average functional connectivity across subjects, with characteristic blocks of correlated regions. The relation between the mean SC and FC matrices is shown in Fig. 4A3 (average correlation 0.50, for extended analysis see SI Table 1) and is similar to other studies [Straathof et al., 2019]. Of course, the observed SC-FC relation may differ depending on the specific pipelines used to estimate the SC and FC. For example, with more conservative preprocessing that aims to suppress potential artifact sources, the individual FC matrices resemble more the typical FC matrix [Kopal et al., 2020] and, in our case, leads to globally decreasing strength of the functional connectivity (for the effects of commonly used preprocessing components see [Bartoň et al., 2019]); at the same time different fiber tracking methods would also lead to varying estimates of SC [Schirner et al., 2015]. SC-FC correlations separately for weakly and strongly connected pairs is in SI Table 1.

**Figure 4:**
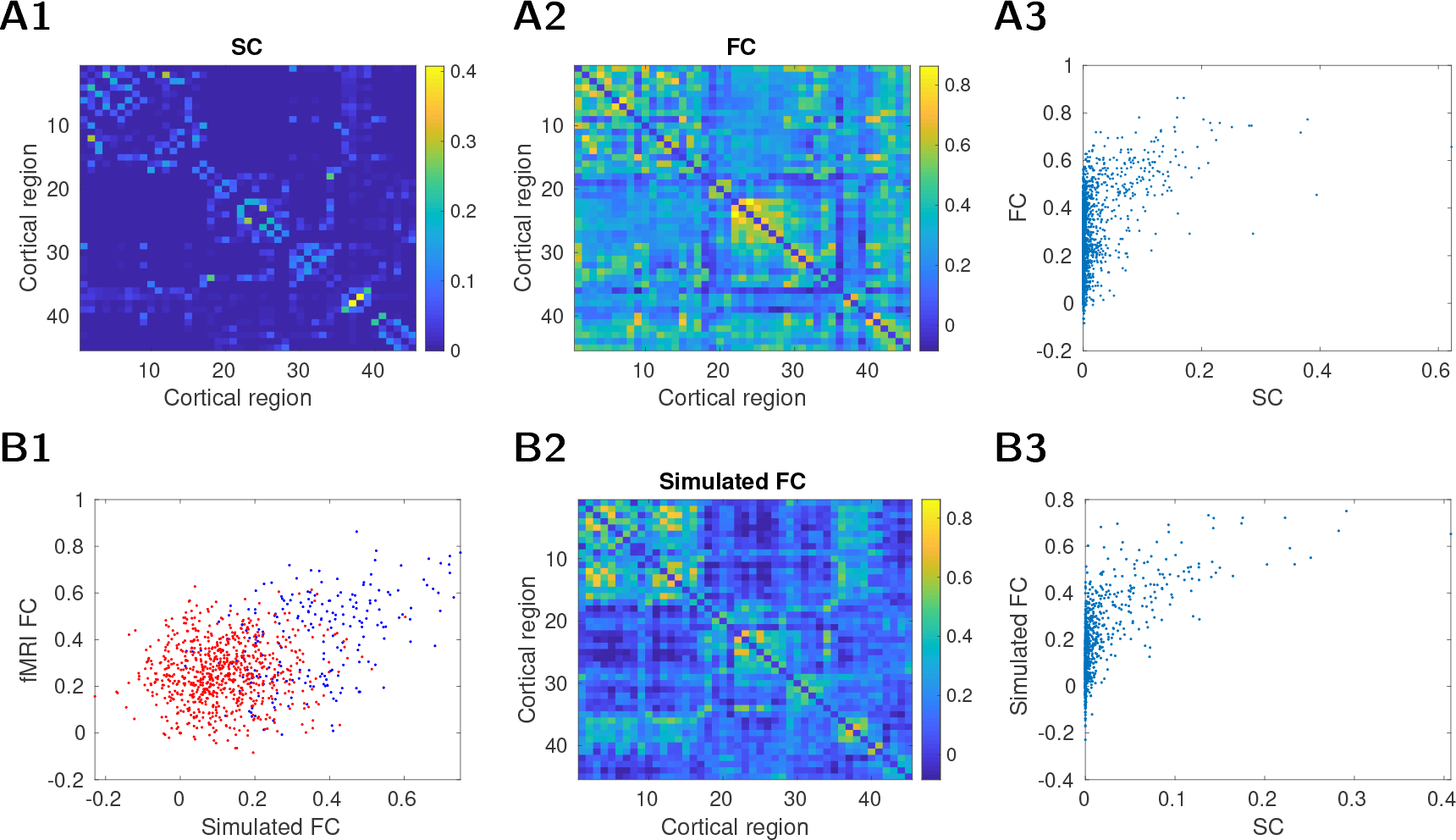
Top. Experimental human dataset. A1. Structural connectivity (SC), an average of 90 control subjects. The strength has an arbitrary scale. The mapping between numbers and their localization is in SI Table 2. A2. Average functional connectivity (FC, 90 subjects). A3. SC-FC relationship avg. correlation was 0.50. Bottom. Model of resting state on the human connectome. B1. Relationship between experimental fMRI FC and FC of the modeled human connectome. The stronger connections (strength > 0.01, see SC above) have blue color. Avg. correlation 0.41 (stronger subset corr. 0.45, weaker subset corr 0.1). B2. FC of the model. B3. SC-FC relationship in the model, avg. correlation 0.63.

The BOLD activity from the computational model of the human connectome largely reproduced the functional blocks of FC (Fig. 4 A2, B2) and the SC/FC relationship (Fig. 4 A3, B3). We then examined how the model’s FC compared with the experimental FC in strong and weak structural connections. Fig. 4B1 shows the relationship between fMRI and the model’s FC pairs - divided into two sets with strong (blue) and weak (red) structural connectivity. The separation in red versus blue points in this plot suggests that the model is able to capture the FC of strongly connected nodes better than weakly connected nodes (the visualization of poorly performing ones on the FC matrix is shown in SI Fig. 11). The SC-FC correlation in our model is similar to other simulation studies [Messé et al., 2015]. Overall, our resting-state model using human connectome had the essential features often observed in human experiments and previous models.

### 2.3 Cholinergic modulation influences global network properties

We next simulated the cholinergic modulation for the human connectome. Similar to rodent DMLN simulation cholinergic modulation was implemented by modifying K^+^leak currents and excitatory AMPA currents. Cholinergic modulation was first applied to all areas equally to simulate a broad ACh release across the brain (for more realistic case when ACh is not released uniformly across all regions see later sections). In this brain-wide high ACh condition, the amplitude of the resting-state activity measured by fALFF and the functional connectivity between regions decreased on average (Fig 5A-B). The reduction in fALFF, as well as in overall FC strength is consistent with rodent DMLN simulations in the previous section. In contrast, the SC-FC correlation increased with ACh (Fig. 5C, panel D shows all SC-FC pairs). This increase was largely driven by a large reduction of FC in low SC ROI-pairs compared to high SC ROI-pairs and is in line with expectations based on a linear approximation of the brain dynamics (see Discussion where we corroborate on this point).

**Figure 5:**
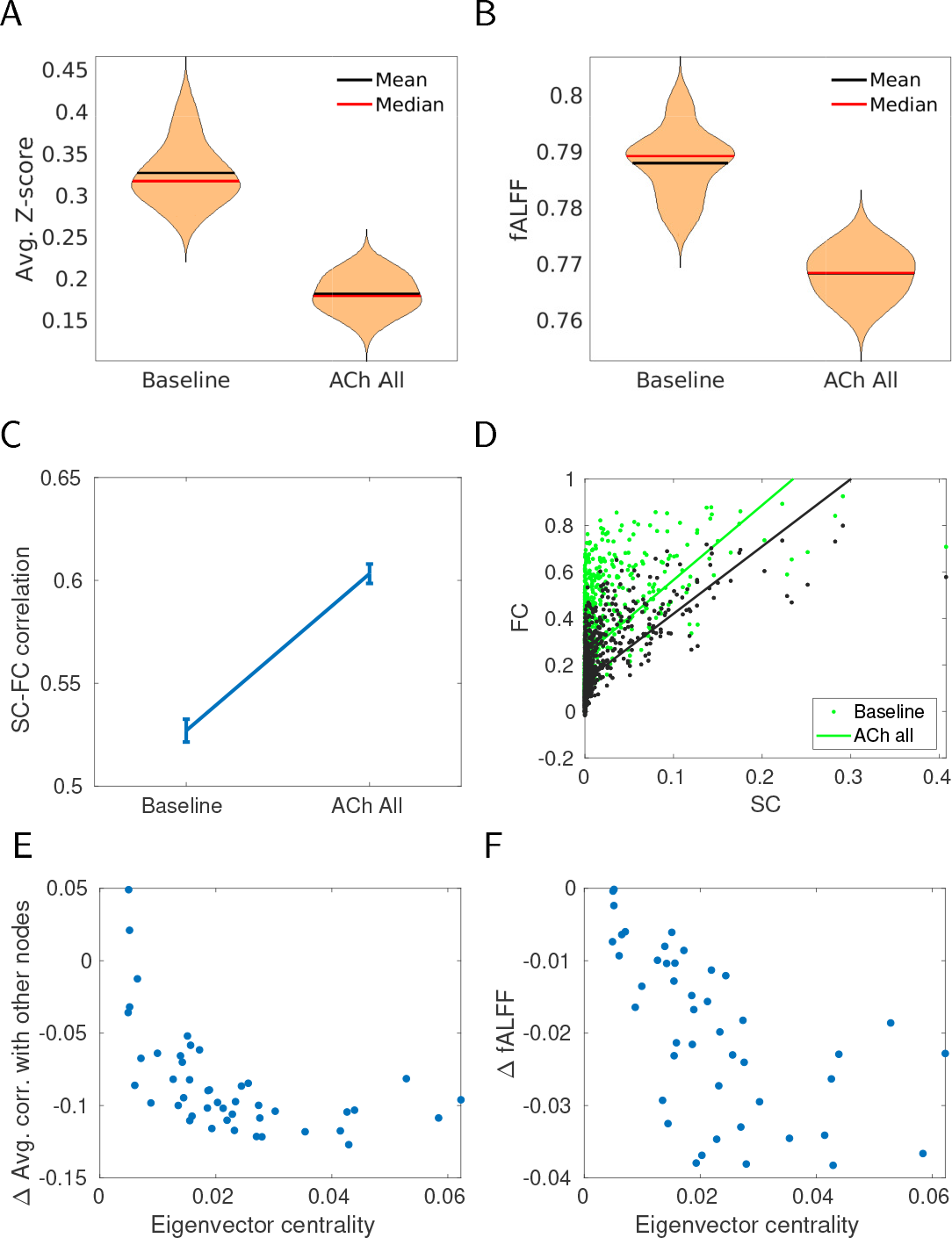
Effect of generic cholinergic release on the connectivity (each condition 20 trials). A. Average functional connectivity of all pairs of nodes for baseline and cholinergic modulation (Z-scored, 20 trials). B. Average fALFF of all nodes for baseline and cholinergic modulation (20 trials). C. Average SC-FC correlation for baseline and cholinergic modulation, 20 trials *±*SEM. D. SC-FC relation for each pair of nodes in the two conditions. E. Decrease of correlation with respect to node centrality. Each point represents one node, the x-axis corresponds to its eigenvector centrality of the node derived from the graph of structural connectivity. y-axis shows decrease of avg. correlation to all other nodes (for a given node). F. Decrease in amplitude (represented by fALFF) for each node of the network. x-axis as in E.

A natural question arises concerning which brain regions should be most affected by the ACh modulation. For any given region there are two contributing factors that determine its response to cholinergic modulation. First, there is the reduction of resting-state activity fluctuations due to changes to excitability and recurrent connections. Second, the input to the region also changes from similar changes in other regions which are projecting to this region. In a network, these two factors interact leading to a cascading effect, with the strongest consequences for the highly connected regions receiving connections from other (highly connected) regions. This notion of overall network-propagated connectivity is conveniently captured in the graph theoretical measure of eigenvector centrality [Bonacich, 1972] well studied in terms of resting-state activity [Lohmann et al., 2010].

We indeed observed that while the cholinergic modulation was applied uniformly, the nodes with higher eigenvector centrality had a larger reduction in resting-state amplitude (Fig 5F). As a consequence, these nodes also had lower FC for its connections. This suggests that the impact of cholinergic modulation depends on the connectivity, with central nodes being more impacted (and some of them residing in DMN). This selectivity further explains the impact of cholinergic modulation of SC-FC relationship.

### 2.4 Differential impact of cholinergic modulation on DMN

We wanted to examine how cholinergic modulation impacts DMN sub-network which has been previously identified in resting-state studies. Previous work from our group [Peeters et al., 2020] and others [Tanabe et al., 2011] suggests a larger influence of cholinergic modulation in DMN compared to task-positive network. One possibility is that DMN is more sensitive to cholinergic modulation, since regions of DMN are the major targets of cholinergic and non-cholinergeric projections of BF [Chandler et al., 2013, Bloem et al., 2014]. In order to examine the selectivity of DMN, we examined the change in resting-state activity and FC under two different conditions: first, whole brain homogeneous ACh release, and second, ACh released only in the DMN areas of the brain (“DMN-only condition”).

Fig. 6A1 shows the average effect on FC in both conditions compared to the baseline. In both brain-wide and DMN-only conditions, there was a significant reduction of FC on average. The steepest decline is visible for the DMN nodes in the DMN-only condition. ACh increase in the entire brain resulted in a larger reduction of FC in DMN compared to DMN only condition and suggests that FC within DMN nodes is sensitive to changes in the rest of the brain.

**Figure 6:**
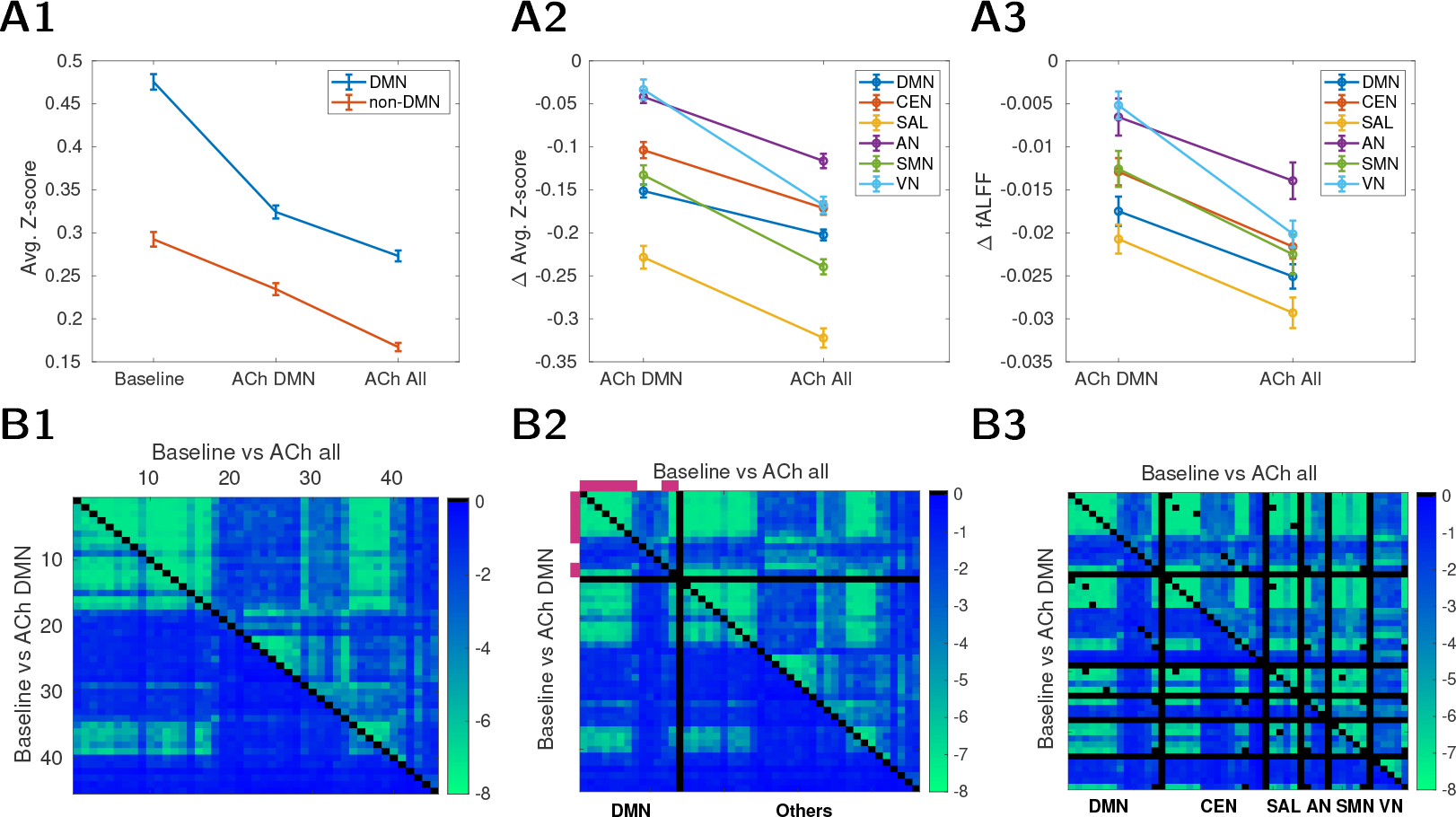
Cholinergic modulation of the human connectome. Three conditions (20 trials for each) are considered: baseline, ACh release only in DMN areas (“ACh DMN”), and ACh release in all brain areas (“ACh all”). A1: Average FC (Fisher z-transformed) computed separately within DMN (blue) and within the remainder of the areas (orange). A2: Changes in FC compared to baseline, projected to major functional networks (CEN=central executive, SAL=salience, AN=auditory, SMN=sensimotor, VN=visual network, for delineations, see SI Table 3). A3: Changes in fALFFs compared to baseline, projected to major functional networks. B: FC changes after ACh release in all (top triangle) / DMN areas (bottom triangle). The color shows log_10_(p-value) of a two-sided Wilcoxon rank sum test, testing the null hypothesis that FC values in baseline and ACh condition are sampled from continuous distributions with equal medians (intuitively, the “blue regions” designate functional connectivities which are not substantially affected by the ACh modulations). Black color is used just as a separator. B1: Areas sorted as in AAL. B2: Areas regrouped to contain DMN and the rest of the areas separately. Violet color indicates DMN regions with higher (in upper 50% regions) eigenvector-centrality index. B3: Areas regrouped by their affiliation to different functional networks. Some nodes are shared across different networks (thus, some self-reference black dots out of the diagonal). For the B3 version without DMN nodes shared in other networks see SI Fig. 13 A2/B2.

We observed several intriguing findings when ACh was increased only in DMN. First, the largest change in FC was observed in the salience network (both within-FC and amplitude were reduced, Fig. 6A2, A3). This is expected as the saliency network shares 50% of nodes with DMN in AAL parcellation (see SI Table 3 and also compare SAL region of Fig. 6B3 bottom vs. SI Fig. 13A2 bottom). When DMN nodes were excluded from the salience network, changes in the rest of the saliency network became less prominent (see SI Fig. 12). More importantly, the sensory regions in auditory and visual networks were not impacted in the DMN-only condition. These findings demonstrate that selective cholinergic modulation of DMN has brain-wide changes in resting-state activity, with the notable exception of the sensory networks. Thus a selective cholinergic modulation of DMN would support the switch between externally oriented task states to internally oriented states (e.g. DMN suppression would not cascade into suppression of the sensory networks).

We then examined the differences in the fine structure of FC across different conditions. Figures 6B show the p-value for each pair of nodes when comparing different conditions. We can see that in the case of whole brain ACh release (top triangle of the matrix), the change of FC generally follows block patterns of hubs seen in FC itself (compare 6B1 top and 4B2), while DMN-only release modulates restricted part of the network (Fig. 6B1 bottom triangle). Reordering the nodes in the matrix so that DMN nodes start first (thus DMN connectivity pairs form left top square matrix, Fig. 6B2), we see that the DMN-only condition causes less global changes but was not limited to DMN regions. DMN itself can be roughly divided into two subsets of nodes – high and low responders to ACh change. We then used the eigenvector centrality as defined in the previous section and found that the nodes with high eigenvector-centrality index (“high influencers” indicated by violet color) had the largest change. The same can be stated as a general rule - high influencers show more ACh-related FC changes in the whole connectome when ACh targets all brain areas, see SI Fig. 13 A1/B1 upper triangle, where we sorted the nodes by their eigenvector-centrality rank.

To offer a comparable analysis to that shown for the rat experiments, we plot the effects of targeting only DMN by ACh in human connectome in a similar vein as in Fig. 3. Again, we observed results matching the experimentally reported pattern [Peeters et al., 2020], namely that DMN nodes under the ACh influence decreased the amplitude in the BOLD signal (Fig. 7A,D), spectrum (Fig. 7C) and functional connectivity (Fig. 7B).

**Figure 7:**
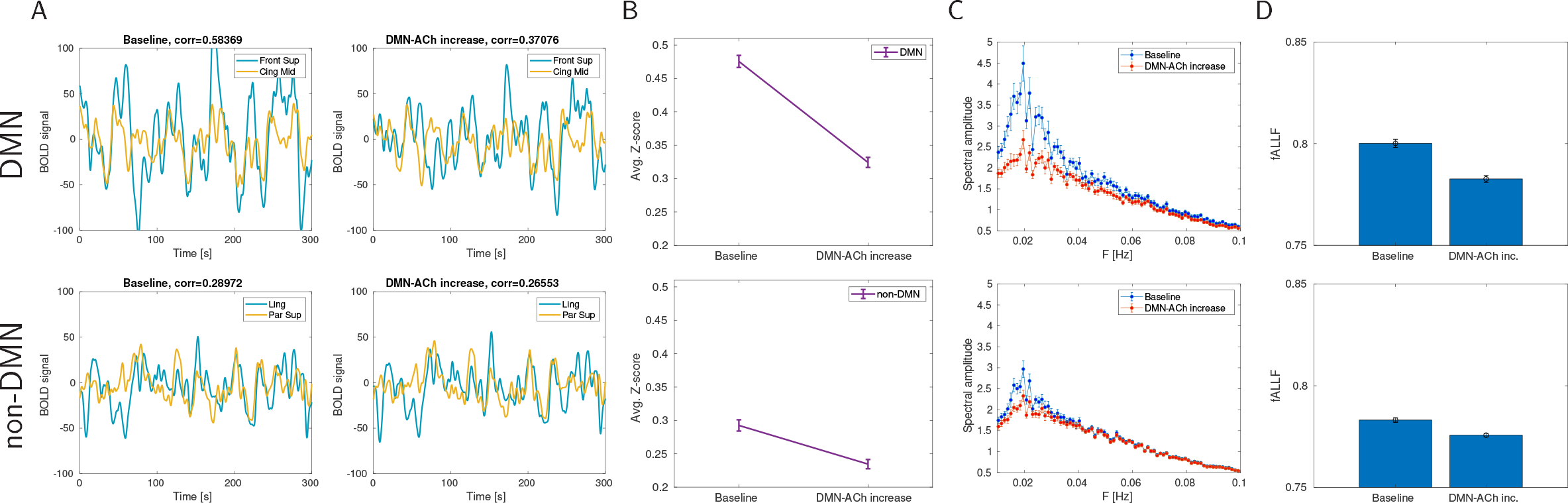
The effect of ACh release in DMN on DMN and non-DMN areas of human connectome. A. Example traces of BOLD signal. Top. Example of two areas in DMN - dorsolateral superior frontal gyrus and median cingulate/paracingulate gyri. Bottom: Example of two areas out of DMN - lingual gyrus and superior parietal gyrus. Left: Spontaneous resting activity of the model (“baseline”), right: the same trial when modulated by ACh release. B. Average functional connectivity comparing baseline and ACh condition (20 trials in each condition), Fisher z-transformed correlation *±*SEM. Top: FC between DMN areas. Bottom: FC between non-DMN areas. C. Average spectral amplitude (*±*SEM, 20 trials) of all areas in DMN (top) and non-DMN areas (bottom) in resting condition (blue) and DMN-modulated-by-ACh condition (red). D. Average fALFF (*±*SEM, 20 trials) of all areas in DMN (top) and non-DMN areas (bottom).

### 2.5 Combined modulation of excitability and excitatory connections explain the state dependent change in resting-state activity

In order to better isolate the neural mechanism of cholinergic modulation of the resting state, we systematically varied the maximal conductance of K-leak current in excitatory neurons and excitatory AMPA connections in a rat DMLN connectome (similar results were observed with human DMN). The baseline (or 100%) condition corresponds to the baseline (resting state) used in rat DMLN simulations. When the K-leak current conductance was reduced (from 110 to 85%), the mean firing rate across neurons increased from silence to 16 Hz (Fig. 8A left and 8B left). In contrast, AMPA conductance had a smaller impact on the firing rate, with the firing rate increasing moderately (3 - 4 Hz variation for lowest K^+^leak conductance). The higher sensitivity of K-leak conductance on firing rate is partly due to the impact of K-leak current on both direct excitability and the indirect effect through its influence on extracellular K^+^concentration. An increase in excitability due to the reduction of K-leak conductance also increases extracellular K^+^ concentration (due to the accumulation of K^+^ions from spikes), which further increases the excitability. This feedback interaction, as reported in our previous models [Fröhlich et al., 2006, 2010, Krishnan and Bazhenov, 2011], may lead to a large non-linear change in firing rate with a change in K-leak conductance.

**Figure 8:**
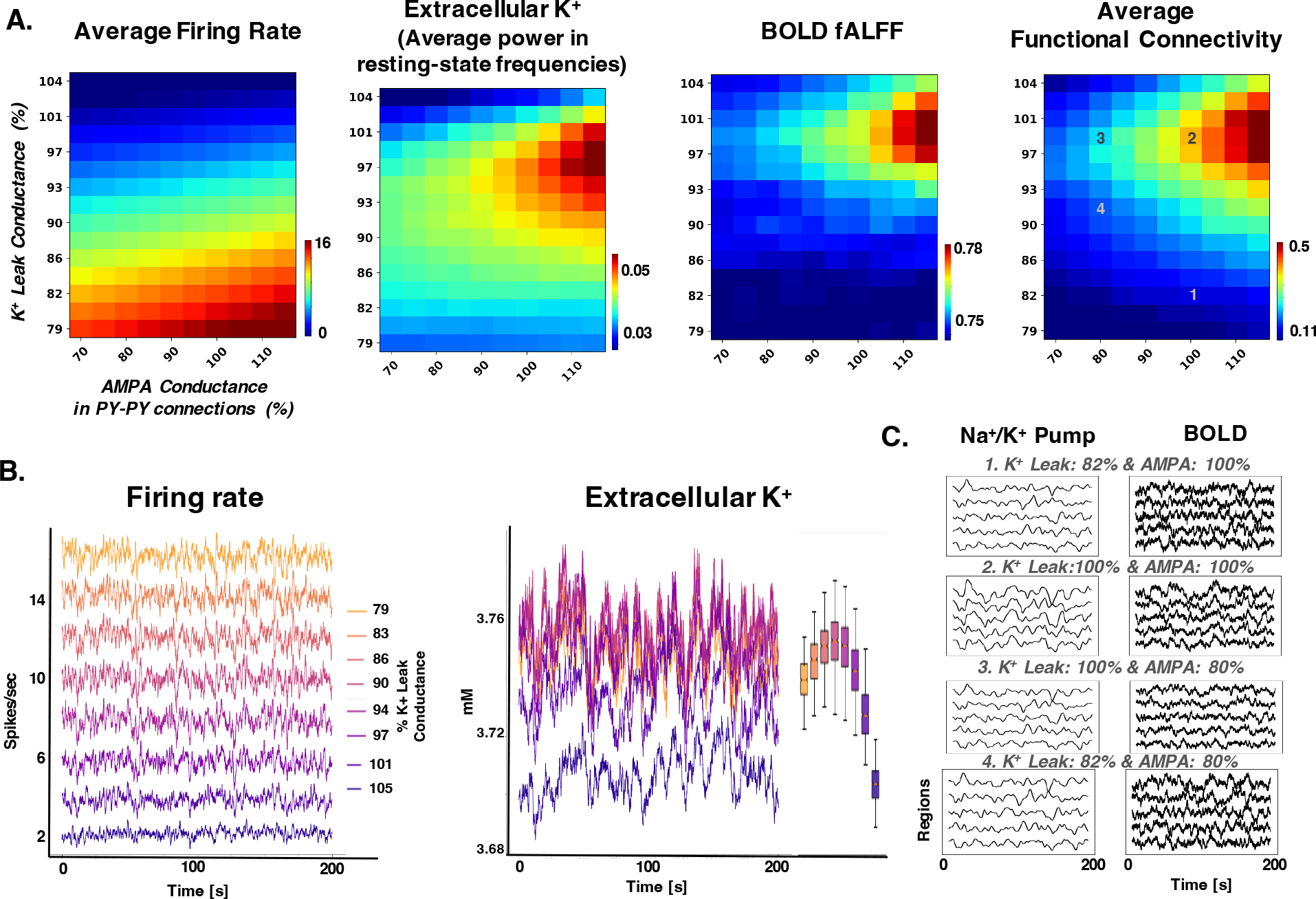
Cellular mechanisms of cholinergic modulation of resting-state activity. A. Different measures of resting-state activity and ion concentration dynamics when K^+^ leak and AMPA conductances are varied. The average firing rate is measured by averaging the mean firing rate across neurons and regions. Extracellular K^+^concentration fluctuation is measured by taking the FFT of the average extracellular K^+^time series. BOLD fALFF, measured from the synaptic activity with fALFF similar to the method described in previous figures. Average functional connectivity is measured as the mean of functional connectivity across all pairs of regions. B. Time series of firing rate and extracellular K^+^for different values of K^+^leak conductance in 100% AMPA condition. The legend shows the correspondence between the color and value of the K^+^leak conductance value for both plots. Inset on the right of the extracellular K^+^ plot is the boxplot with whiskers corresponding to the 1 and 99 percentile of the data to indicate the range of the extracellular K^+^fluctuations. C. Na^+^/ K^+^pump and BOLD time series for different regions for selected conditions. The number in the title corresponds to the number in the 2D sweep image in the right panel in A.

In contrast to the firing rate, the fluctuations in extracellular K^+^concentration and BOLD in resting-state frequencies were impacted by both K-leak and AMPA conductances. Its value increased with AMPA conductance only for the intermediate values (around 100%) of K-leak conductance. Specifically, the extracellular K^+^concentration doubled its value with AMPA conductance changed from 70 to 100% only for the intermediate values of K-leak conductance. The extracellular K^+^concentration fluctuation is lower at high K-leak conductance (above 102%) due to the significant drop in the excitability and the overall quiescence in the network activity. Interestingly, at low K-leak conductance values, when the firing rate is high, there is also the reduction in K^+^fluctuations due to smaller contribution of K^+^ions from K-leak currents and lower synchronization between regions (as shown by lower functional connectivity Fig. 8A).

The BOLD, average functional connectivity (Fig. 8A), and Na^+^/ K^+^pump activity (Fig. 8C) closely matched the trend in the extracellular K^+^ concentration. Both BOLD and Na^+^/ K^+^pump had the highest values for the intermediate values of K^+^leak and AMPA conductance. Further, the synchronization in BOLD (Fig. 8A right) and Na^+^/ K^+^pump (Fig. 8C) increased with AMPA conductance only in the intermediate range of K^+^leak conductance. The reduction in recurrent excitation led to lower BOLD, functional connectivity mediated by the changes to extracellular K^+^fluctuations (Fig. 8C, panel 3). Thus, these findings suggest that the interaction between the excitability of neurons and the strength of recurrent excitation critically determines the fluctuations of K^+^in resting-state activity, Na^+^/ K^+^pump, and the BOLD. Further, our findings suggest that cholinergic modulation could also result in any of the intermediate values of K-leak and AMPA conductance, allowing for gradual and selective changes in resting-state activity.

Thus, cholinergic modulation which influences both excitability and recurrent excitation is ideally suited for modulating resting-state activity and its functional connectivity.

## 3 Discussion

The widely used “resting-state” activity is thought to reflect intrinsic spontaneous activity when the brain is in rest periods [Raichle, 2015]. While the general spatiotemporal patterns of rest and task-related activity are surprisingly similar [Smith et al., 2009], some sub-networks, in particular the default mode network, have increased activity during rest periods as compared to a range of cognitive tasks [Shulman et al., 1997, Raichle and Snyder, 2007]. Moreover, during sustained demanding cognitive tasks, DMN activity fluctuation, while still apparent, is substantially reduced compared to the DMN activity fluctuation during an unconstrained resting state [Fransson, 2006, Northoff et al., 2010]. In this study, we first tested the hypothesis that cholinergic activation promotes the inhibition of resting-state activity in DMN. To develop our model we used changes in rsfMRI following direct cholinergic neuron stimulation in rodents. We then extend the model to human connectome derived from DW-MRI measurements obtained from healthy human subjects. Results from our study support the hypothesis that a change in neuromodulation supports the switching between different functional networks.

Neuromodulation plays a critical role in the switch between different states of vigilance [Lee and Dan, 2012]. In addition, neuromodulation is also proposed to play an essential role in shaping the dynamics of the task-and resting-state networks [Thiele and Bellgrove, 2018, Shine, 2019, van den Brink et al., 2019]. Acetylcholine is a major neuromodulator released through broad projections from the basal forebrain [Picciotto et al., 2012]. We used data from a previous experiment [Peeters et al., 2020], which involved selective activation of the cholinergic neurons in the basal forebrain using excitatory DREADDs. Simultaneously measuring resting-state fMRI with DREADDs showed suppression of rsfMRI during cholinergic neuron activation.

The results from the experiments were then used to constrain the computational model of DMLN with the connectivity based on NeuroVIISAS atlas for rodent brain [Schmitt and Eipert, 2012]. We used a computational model of resting-state activity that included realistic ionic and synaptic currents based on the Hodgkin-Huxley formulation and ion dynamics. In this model, the interaction between cellular currents and ion dynamics leads to slow resting-state activity [Krishnan et al., 2018]. Using a biophysical model allowed us to implement cholinergic activation as a reduction of conductance of K^+^leak current and excitatory connections. The cellular action of acetylcholine is not fully understood and could be variable across neurons [Colangelo et al., 2019]. Thus, in this work we examined the most prominent action of cholinergic modulation involving K^+^leak and excitatory connections. We also examined a more elaborate model of cholinergic modulation involving all K^+^, AMPA and NMDA currents, which resulted qualitatively similar to model involving only changes to K^+^ leak and AMPA currents. The model was able to replicate the reduction in resting-state activity and its functional coupling following cholinergic activation.

An increase in DMN activity is a hallmark of internally oriented states and the transition between those states and states with externally oriented attention is poorly understood [Buckner and DiNicola, 2019]. Several – not necessarily exclusive – candidates for the control of the transition are the activity of other functional networks [Sridharan et al., 2008, Menon and Uddin, 2010, Spreng et al., 2013], thalamocortical circuits, and ACh-dependent pathway mediated by basal forebrain (BF) [Nair et al., 2018, Buckner and DiNicola, 2019], which can be (together with the thalamus) thought of as a subcortical part of DMN [Alves et al., 2019]. Our results suggests that selective cholinergic modulation of DMN could facilitate this switch.

While it is known that BF projects broadly over the neocortex and the projections can be very specific in mammals [Gratwicke et al., 2013, Bloem et al., 2014, Zaborszky et al., 2015a, Záborszky et al., 2018, Markello et al., 2018], it is notoriously difficult to get exact human connectivity due to the small volume of the critical areas [Zaborszky et al., 2008, Chiang-shan et al., 2014]. We opt for the hypothesis that important DMN regions are more affected by cholinergic release (due to either higher probability in direct projections [Nazari et al., 2022], or by the graded density of AChR receptors, or by a partial activity of specific regions of nucleus basalis of Meynert translating into partial neocortical activations). This is, of course, not the only possibility and there are more complicated accounts of how ACh acts globally [Turchi et al., 2018] and within BF circuits [Yang et al., 2017, Gielow and Zaborszky, 2017, Espinosa et al., 2019a,b]. As human experiments are limited to broad nicotine-related manipulations we initially show an animal model where we could directly influence major cholinergic center (BF), which in turn affects core hubs of DMN – or more precisely, their animal DMLN analogues which were shown to be present across many species [Vincent et al., 2007, Rilling et al., 2007, Popa et al., 2009, Lu et al., 2012, Stafford et al., 2014, Zhou et al., 2016].

In this study, we only examined the cholinergic modulation arising from the basal forebrain. However, major projections of BF includes glutamatergic and GABAergic projections, which were not examined here. It has been proposed that glutamatergic and GABAergic projections mediate the BF influence on cortex [Kim et al., 2015] and in particular DMN [Nair et al., 2018, Lozano-Montes et al., 2020]. Specifically, there is an increase in glutamatergic input and a reduction of GABAergic input to cortex when the cholinergic neuron is not active [Yang et al., 2014]. Such an increase in glutamatergic input would further increase resting-state activity during rest periods in our model, and the overall results will be qualitatively consistent with the results reported in the current study. A more detailed computational model of BF subtypes could, in the future, isolate the cell type specific mechanisms.

Our model of resting state with human connectome captured several features of rsfMRI in humans, including the SC-FC relationship [Honey et al., 2009, Hermundstad et al., 2013]. The relation between SC and FC is far from straightforward, and while they are correlated, there is no simple match. Instead, there is a significant correlation variability reported across the studies [Straathof et al., 2019]. However, our aim here was not to optimize SC-FC correspondence. Firstly, this was already attempted in multiple simulation studies [Messé et al., 2015]; moreover, optimizing solely for the best match can even be detrimental to the models’ dynamical properties [Hansen et al., 2015]. Our model, however, shows a quantitative correspondence similar to the reported results. The functional connections directly supported by existing structural/anatomical connections were captured well, while weak structural connections rendered the prediction of functional coupling weak. The SC-FC match between the model and data could be improved if specific measures supporting cytoarchitectonic, transcriptomic, and higher order interactions were added to the model [Suárez et al., 2020]. However, despite these additions, there remains a fundamental problem: human brain tractography is inherently limited and does not capture gray matter tracts and fibers going through thick bundles of axons, e.g. corpus callosum [Thomas et al., 2014, Maier-Hein et al., 2017]. Hence SC typically misses interhemispheric connections known to affect FC [Messé et al., 2014, Roland et al., 2017]. Another problem stems from the fact that experimental FC values depend on specific parameters for preprocessing pipeline of the BOLD signal. Thus the predictive power of our model might not be directly comparable to other studies using different experimental SC/FC datasets.

We also observed that the SC-FC match increased with an increase in ACh (which generally reduces coupling between the nodes). This is in line with the expectations based on a simple linear approximation of the brain dynamics. In a linear process with weak coupling, the correlation matrix of the time series basically copies the structure of the coupling matrix, as only the first order interactions give rise to correlations of sufficient strength to be above the noise level. However, for a system with stronger coupling, the correlations due to indirect links (such as due to common source(s), or multiple steps of a causal chain) become strong enough to cause correlations (functional connectivity) above the noise level [Liégeois et al., 2020]. Thus, for strongly coupled systems, the functional connectivity can deviate further from the structural connectivity.

In human connectome simulations, we concentrated on a particular role of ACh in switching the balance in the neocortical state from default mode network activation to activation of networks involved in external sensory processing [Tanabe et al., 2011, Klaassens et al., 2017] or executive control [Sutherland et al., 2015]. DMN, a network active mainly in the resting condition [Greicius et al., 2003], was associated with a variety of internally oriented mental states [Andrews-Hanna et al., 2014, Konishi et al., 2015] while inhibited with external goal-oriented tasks [Fox et al., 2005, 2009, Andrews-Hanna et al., 2014]. Thus, we considered global cholinergic modulation of the entire brain and selective cholinergic modulation only in DMN. Cholinergic modulation only in DMN regions translated to a picture consistent with the experimental results in rodents [Peeters et al., 2020]; moreover, it showed DMN-specific inhibition which did not appear when ACh was uniformly affecting all cortical regions. Within the DMN, the modulation mainly affected the areas with higher eigenvector-centrality rank (i.e., highly connected nodes preferring connections to other highly connected nodes). As DMN richly connects (and even shares some nodes) with other functional networks, ACh-triggered changes in DMN propagate to other parts of the brain, however to a lesser degree than it is the case of uniform cholinergic release across all the areas; notably, the change did not impact internal coupling within visual/auditory networks. This observation makes cholinergic-related suppression of DMN compatible with independent activity in sensory regions connected with attention to external stimuli.

In conclusion our findings suggest cholinergic modulation on a cellular level leads to changes in large scale dynamics powerful enough to be a vital part of the intrinsic switching mechanism between different brain networks.

## 4 Materials and Methods

### 4.1 Biophysical model

The microcircuit connectivity and dynamics is identical to our previous work [Krishnan et al., 2018]. To briefly summarize, each network area (ROI) consists of 50 excitatory and 5 inhibitory neurons. Both excitatory and inhibitory neurons were modeled via axosomatic and dendritic conductance-based compartments following the equations

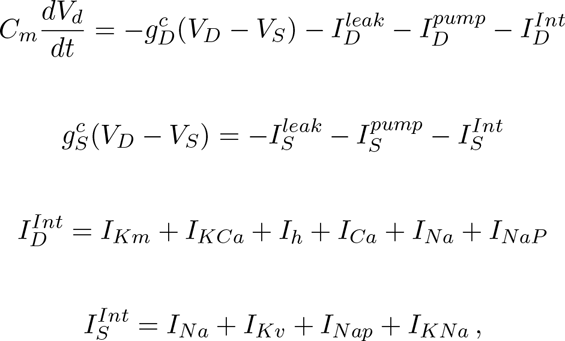

where *C_m_* is membrane capacitance, *V_D,S_* are dendritic/axosomatic compartment voltages, 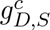 are leakage conductances, 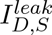 are sums of the ionic leak currents, 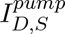 are sums of Na^+^ and K^+^ currents through Na^+^/K^+^ pump, 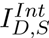 are intrinsic currents. The dendritic compartment includes fast sodium current (*I_Na_*), persistent sodium current (*I_NaP_*), slowly activating potassium current (*I_Km_*), calcium-activated potassium current (*I_KCa_*), hyperpolarization-activated depolarizing mix cationic currents (*I_h_*), high threshold Ca^2+^current (*I_Ca_*) and leak currents [Krishnan and Bazhenov, 2011, González et al., 2015, Krishnan et al., 2015]. The axosomatic compartment includes fast sodium current (*I_Na_*), persistent sodium current (*I_NaP_*), delayed-rectifier potassium current (*I_Kv_*) and sodium-activated potassium current (*I_KNa_*). Ion concentrations were modeled for intracellular K^+^, Na^+^, Cl*^−^*,Ca^2+^ and extracellular K^+^, Na^+^. K^+^/Na^+^ pump for K^+^/Na^+^ regulation and KCC2 cotransporter for extrusion of Cl*^−^* were used for both neuron types [Bazhenov et al., 2004, Fröhlich and Bazhenov, 2006, Krishnan and Bazhenov, 2011, González et al., 2015].

Extracellular space was modeled for each neuron with local ion diffusion between nearest neighbors. It was tightly bounded between the glia and neurons, and there was an instantaneous and direct impact of ion concentration changes in the extracellular space on the neuronal and glial activity. Glial regulation of extracellular K^+^ was modeled as a free buffer [Krishnan and Bazhenov, 2011, Krishnan et al., 2015, González et al., 2015].

Local connectivity within single cluster was mediated via AMPA/NMDA conductances for PY->PY/IN and GABA_A_ for IN->PY, local connectivity radiuses were *r*(Py *→* Py) *≤* 5, *r*(In *→* In) *≤* 2, *r*(In *→* Py) *≤* 5*, r*(Py *→* In) *≤* 1. Long range connections between clusters *i → j* was mediated via AMPA conductances with 15% probability of Py *∈ i* connecting to Py *∈ j*, and GABA_A_ conductances with restricted convergence of 5 Py *∈ i* to 1 In *∈ j* and probability of connection 25% for each possible connection.

Structural connectivity of human connectome was used (90 ROIs from AAL template, 45 for a single hemisphere, acquisition is described below), for modulation of DMN the subset of areas participating in default mode was defined by Lee and Frangou [2017] (see SI Table 3 for explicit list of the nodes). Structural connectivity of rat DMLN (see Fig. 1A) was extracted from the NeuroVIISAS database of axonal tracing studies [Schmitt and Eipert, 2012].

The BOLD signal was created for each cluster first by averaging total synaptic input for all excitatory neurons in 100 ms windows and then it was convolved with a hemodynamic response function imported from Statistical Parametric Mapping (SPM) package [Ashburner et al., 2014].

### 4.2 Human data - structural connectivity

Acquisition of MRI data and construction of structural connectivity was identical to the methods described in Škoch et al. [2022]. To summarize, the data provided here are based on MRI scans of 90 healthy control individuals participating in the Early-Stage Schizophrenia Outcome study [Melicher et al., 2015]. The study was conducted in accordance with the Declaration of Helsinki. The local Ethics Committee of the Prague Psychiatric Center approved the protocol on 29 June 2011 (protocol code 69/11). The construction of structural connectivity matrices was based on a connectome generated by probabilistic tractography on diffusion MRI data. We used ROIs from the widely used AAL atlas (Automated Anatomical Labeling atlas, [Tzourio-Mazoyer et al., 2002]). The connectivity between two ROIs is based on the number of streamlines in the tractogram beginning in one ROI and terminating in the other ROI. Accurate mapping of the AAL atlas ROIs to the diffusion data space was realized as a two-stage process: affine mapping of structural T1 images to MNI space and a rigid-body mapping between the T1 structural data and the DWI data, both for each subject.

We performed the MRI scanning at the Institute for Clinical and Experimental Medicine in Prague, on a 3 T Trio Siemens scanner (Erlangen, Germany). A 12-channel head coil was used, software version syngo MR B17. DWI data were acquired by a Spin-Echo EPI sequence with TR/TE = 8300/84 ms, matrix 112 *×* 128, voxel size 2 *×* 2 *×* 2 mm3, b-value 0 and 900 s/mm2 in 30 diffusion gradient directions, 2 averages, bandwidth 1502 Hz/pixel, GRAPPA acceleration factor 2 in phase-encoding direction, reference lines 24, prescan normalize off, elliptical filter off, raw filter on – intensity: weak, acquisition time 9:01. T1 3D structural image was acquired by using the magnetization prepared rapid acquisition gradient echo (MPRAGE) sequence with (TI – inversion time) TI/TR/TE = 900/2300/4.63 ms, flip angle 10*°*, 1 average, matrix 256 *×* 256 *×* 224, voxel size 1 *×* 1 *×* 1 mm3, bandwidth 130 Hz/pixel, GRAPPA acceleration factor 2 in phase-encoding direction, reference lines 32, prescan normalize on, elliptical filter on, raw filter off, acquisition time 5:30.

### 4.3 Human data - functional connectivity

#### fMRI data acquisition

Scanning was performed with a 3T MRI scanner (Siemens Magnetom Trio) located at the Institute for Institute of Clinical and Experimental Medicine in Prague, Czech Republic. Functional images were obtained using T2-weighted echo-planar imaging (EPI) with blood oxygenation level-dependent (BOLD) contrast using SENSE imaging. GE-EPIs (TR/TE=2000/30 ms, flip angle=70*°*) comprised of 35 axial slices acquired continuously in sequential decreasing order covering the entire cerebrum (voxel size=3*×*3*×*3 mm, slice dimensions 48×64 voxels). 400 functional volumes were used for the analysis. A three-dimensional high-resolution MPRAGE T1-weighted image (TR/TE=2300/4.63 ms, flip angle 10*°*, voxel size=1*×*1*×*1 mm) covering the entire brain was acquired at the beginning of the scanning session and used for anatomical reference.

#### Data preprocessing, brain parcellation, and FC analysis

The rsfMRI data were corrected for head movement (realignment and regression) and registered to MNI standard stereotactic space (Montreal Neurological Institute, MNI) with a voxel size of 2*×*2*×*2 mm by a 12 parameter affine transform maximizing normalized correlation with a customized EPI template image. This was followed by segmentation of the anatomical images in order to create subject-specific white-matter and CSF masks. Resulting anatomical images and masks were spatially normalized to a standard stereotaxic MNI space with a voxel size of 2*×*2*×*2 mm.

The denoising steps included regression of six head-motion parameters (acquired while performing the correction of head-motion) and the mean signal from the white-matter and cerebrospinal fluid region. Time series from defined regions of interest were additionally filtered by a band-pass filter with cutoff frequencies 0.004 - 0.1 Hz. The regional mean time series were estimated by averaging voxel time series within each of the 90 brain regions (excluding the cerebellar regions) comprising the Automated Anatomical Labeling (AAL) template image [Tzourio-Mazoyer et al., 2002]. To quantify the whole-brain pattern of functional connectivity, we performed a ROI-to-ROI connectivity analysis and computed for each subject the Pearson’s correlation matrix among the regional mean time series, as (linear) Pearson’s correlation coefficient has been shown to be suitable for fMRI ROI functional connectivity estimation [Hlinka et al., 2011].

### 4.4 Animal data

Experimental framework and acquisition of MRI data is identical to [Peeters et al., 2020]. To summarize, 28 adult ChAT-Cre Long Evans rats were used, of which 14 males and 14 females. Animals were group housed with a 12h light/dark cycle and with controlled temperature (20 – 24*°*C) and humidity (40%) conditions. Standard food and water were provided ad libitum. All procedures were in accordance with the guidelines approved by the European Ethics Committee (decree 2010/63/EU) and were approved by the Committee on Animal Care and Use at the University of Antwerp, Belgium (approval number: 2015-50).

All rats received stereotactic surgery targeting the right nucleus basalis of Meynert, horizontal diagonal band of broca and substantia innominata to transfect cholinergic neurons using either a Cre-dependent DREADD virus (AAV8-hSyn-DIO-hM3Dq(Gq)-mCherry) (N = 16) or sham virus (AAV8-hSyn-DIO-mCherry) (N = 12). Resting-state functional MRI was performed at least two months after surgery, to allow stable expression of the virus. Animals were anesthetized using a combination of isoflurane (0.4%) and medetomidine (0.05 mg/kg bolus followed by a 0.1mg/kg/hr continuous infusion). An intravenous catheter was placed in the tail vein which was used to administer 1 mg/kg CNO or saline. A gradient-echo EPI sequence was used (TE: 18 ms, TR: 2000ms, FOV: (30 x 30) mm2, matrix [128 x 96], 16 slices of 0.8 mm) on a 9.4T Bruker Biospec preclinical MRI scanner. A 5 minute baseline scan was followed by the injection of either CNO or saline during a 20 minute scan, followed by another 5 minute rsfMRI scan. DREADD expressing animals received two scan sessions, one with an injection of CNO and second session with an injection of saline, while Sham animals only received one scan session with CNO.

Preprocessing of the rsfMRI data included realignment, spatial normalization to a study specific template, masking, smoothing and filtering (0.01-0.2 Hz) using Matlab 2014a and SPM12 software [Ashburner et al., 2014]. Region-of interest based analysis was performed using predefined regions belonging to the default mode network. Amplitude of low frequency fluctuations were extracted from seed regions (cingulate cortex and retrosplenial cortex) within the right hemisphere. FC and fALFF values were compared before and after injection of CNO/saline using paired two-sample t-tests or unpaired two-sample t-tests.

## Acknowledgements

P.S. and J.H. were supported by the Czech Science Foundation project No. 21-32608S, JH and AS were supported by MH CZ – DRO (NUDZ, 00023752), M.v.d.B. and G.A.K. were supported by the Fund of Scientific Research Flanders (G048917N), AS was supported by MH CZ – DRO (IKEM, IN: 00023001), M.B. and G.K. were supported by National Science Foundation Grant (2209874) and National Institutes of Health grants (1R01MH117155, 1R01MH125557, 1R01NS104368, 1R01NS109553).

Authors have no conflict of interest to declare.

## 5 Figure/table supplements

### 5.1 Comparison of detailed and simplified model of cholinergic modulation

**Figure 9:**
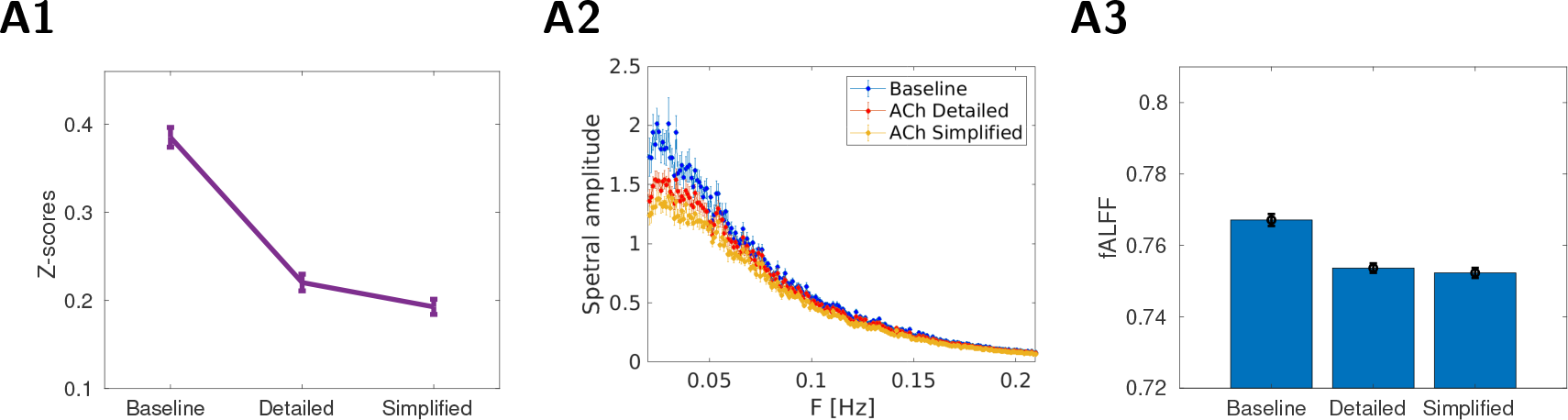
Comparison of detailed model of cholinergic modulation and it’s simplified version. A1: Average functional connectivity comparing baseline and ACh condition for detailed and simplified model (16 trials in each condition), Fisher z-transformed correlation *±*SEM. B2: Average power-spectra from all DMLN areas (*±*SEM, 16 trials). The blue curve is the baseline condition, red/orange ACh modulation in detailed/simplified model. B3: fALFF values calculated from all DMLN areas. The bar graphs present mean fALFF values *±*SEM.

### 5.2 Human connectome

**Figure 10:**
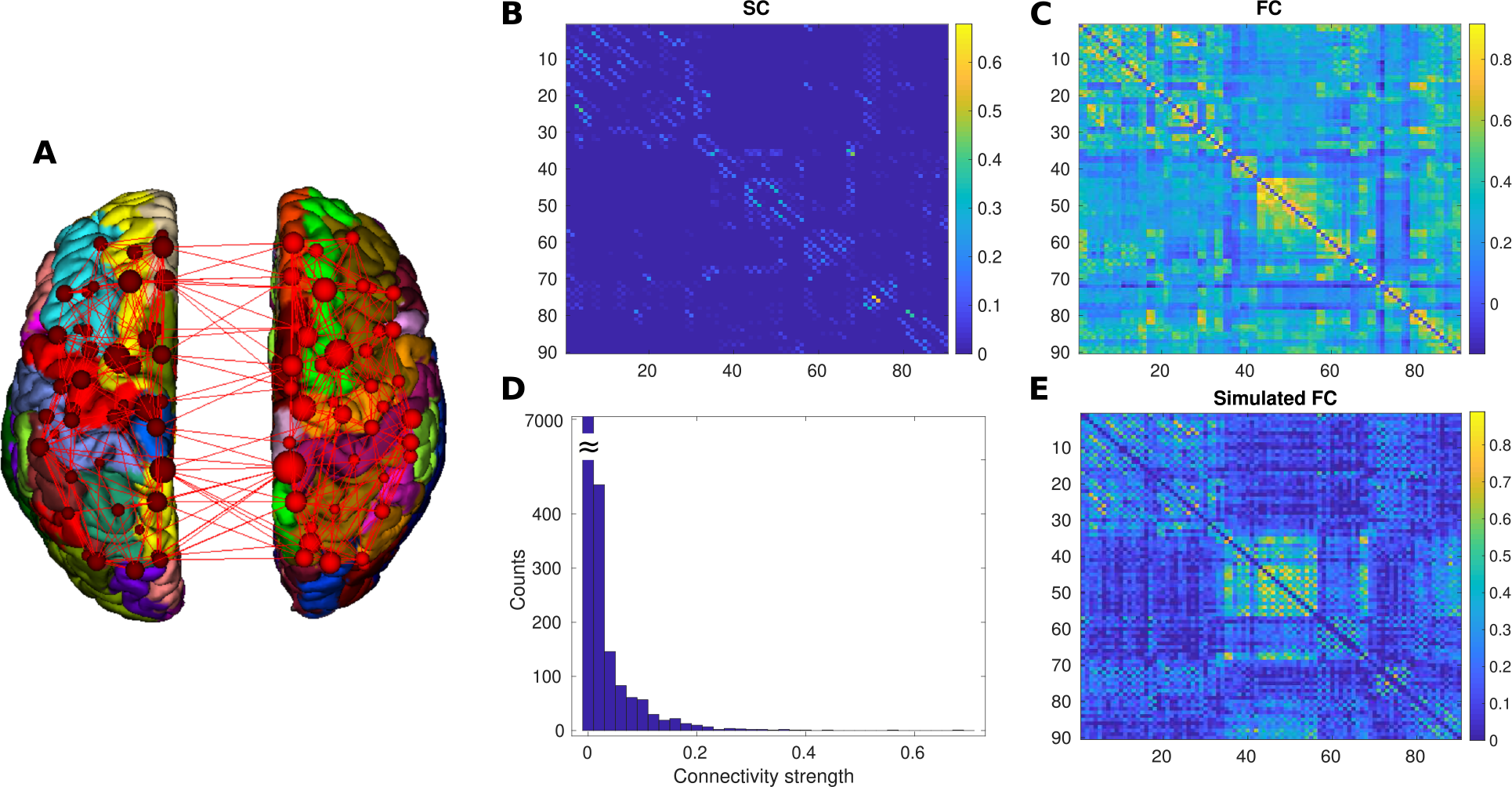
Experimental human dataset. A. Visualization of AAL parcelation. B. Average structural connectivity (Both hemispheres, 90 regions, AAL atlas). Large subset of all underlying SCs can be downloaded from [Škoch et al., 2022]. C. Average functional connectivity from identical 90 subjects. D. Distribution of coupling strengths. E. Functional connectivity obtained by simulation of both hemispheres.

### 5.3 Visualization of poorly performing correlation pairs

**Figure 11:**
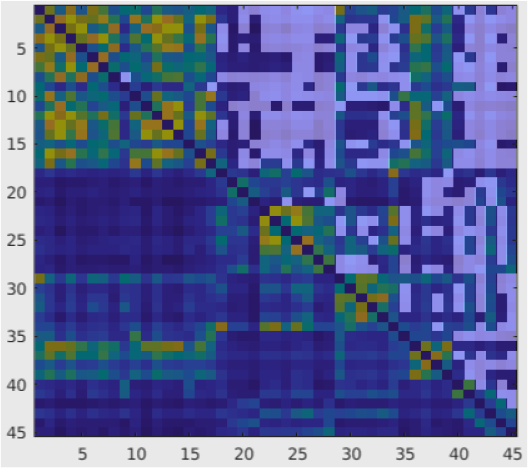
Visualization of pair of nodes (white map) for which the model predicts lower correlation than would be expected from experimental fMRI FC. FC from simulated data used as a background.

### 5.4 Effect of various transformations on average SC/FC values

**Table 1:**
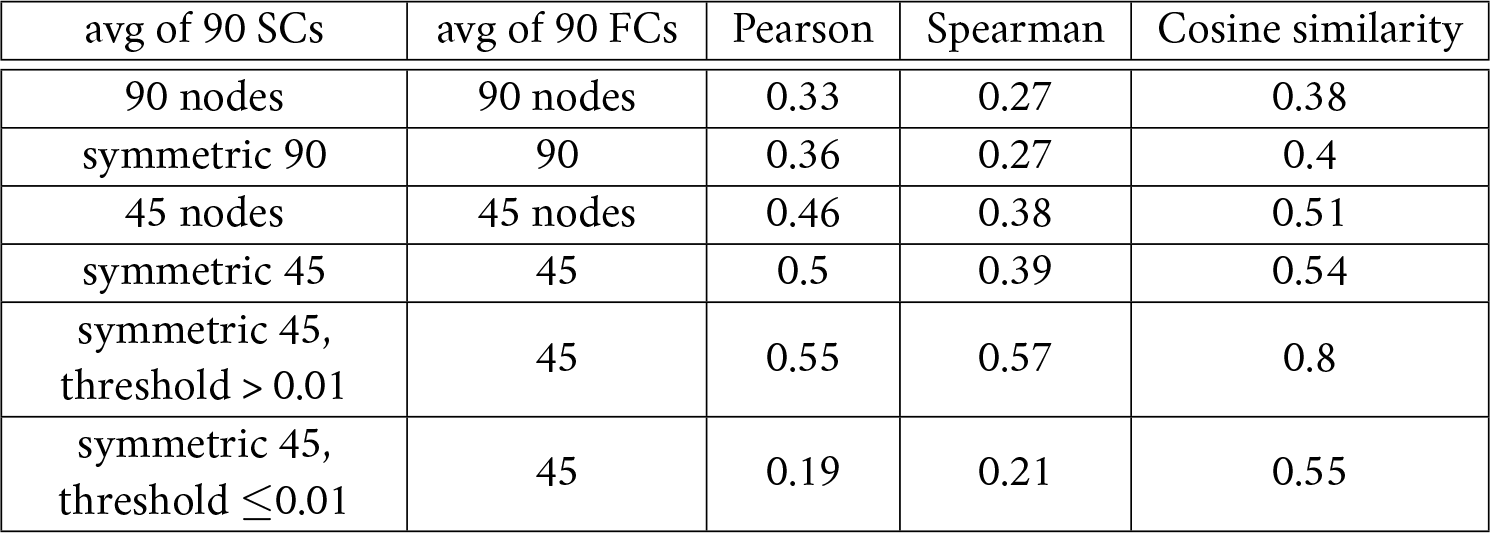
Average correlation of SC/FC matrix for different conditions and transformations. Both hemispheres contain 90 regions. SC is not entirely symmetric, but due to limits of structural imaging the orientation can not be obtained. Thus the symmetric version of connectivity matrix is often considered. The bottom half of the table shows the same comparison for single hemisphere only (45 brain regions). In the last two rows we apply threshold on SC strength, so that either only anatomically “existing” (above threshold 0.01) or “non-existing” (below 0.01) connections are used to compute correlation with their FC counterparts.

### 5.5 AAL parcellation

**Table 2:** Mapping between node numbers and their naming in AAL parcellation scheme for a single hemisphere.

| Index | ROI name | Index | ROI name | Index | ROI name |
| --- | --- | --- | --- | --- | --- |
| 1 | Precentral | 16 | Cingulum Ant | 31 | Parietal Inf |
| 2 | Frontal Sup | 17 | Cingulum Mid | 32 | SupraMarginal |
| 3 | Frontal Sup Orb | 18 | Cingulum Post | 33 | Angular |
| 4 | Frontal Mid | 19 | Hippocampus | 34 | Precuneus |
| 5 | Frontal Mid Orb | 20 | ParaHippocampal | 35 | Paracentral Lobule |
| 6 | Frontal Inf Oper | 21 | Amygdala | 36 | Caudate |
| 7 | Frontal Inf Tri | 22 | Calcarine | 37 | Putamen |
| 8 | Frontal Inf Orb | 23 | Cuneus | 38 | Pallidum |
| 9 | Rolandic Oper | 24 | Lingual | 39 | Thalamus |
| 10 | Supp Motor Area | 25 | Occipital Sup | 40 | Heschl |
| 11 | Olfactory | 26 | Occipital Mid | 41 | Temporal Sup |
| 12 | Frontal Sup Medial | 27 | Occipital Inf | 42 | Temporal Pole Sup |
| 13 | Frontal Mid Orb | 28 | Fusiform | 43 | Temporal Mid |
| 14 | Rectus | 29 | Postcentral | 44 | Temporal Pole Mid |
| 15 | Insula | 30 | Parietal Sup | 45 | Temporal Inf |

**Table 3:** Definitions of functional networks in the AAL template taken from [Lee and Frangou, 2017]. Direct mapping on AAL ids is courtesy of Won Hee Lee and Sophia Frangou. The most-right column is the reduction on a single hemisphere (45 nodes, in case of the few non-symmetric nodes we use the covering set).

| Functional network | Nodes IDs in AAL parcellation | Covering set in a single hemisphere |
| --- | --- | --- |
| DMN<br>(Default mode) | 3 4 5 6 7 8 23 24 25 26 31 32 33 34 35 36<br>37 38 39 40 65 66 67 68 77 78 | 2 3 4 12 13 16 17 18 19 20 33 34 39 |
| CEN<br>(Central executive) | 3 4 7 8 9 14 23 59 61 62 64 65 66 67<br>72 77 85 89 | 2 4 5 7 12 30 31 32 33 34 36 39 43 45 |
| SAL<br>(Saliency) | 7 8 19 20 29 30 31 32 | 4 10 15 16 |
| AN<br>(Auditory) | 78 79 80 81 82 | 39 40 41 |
| SMN<br>(Sensimotor) | 1 2 20 57 58 69 77 78 | 1 10 29 35 39 |
| VN<br>(Visual) | 43 44 49 50 51 52 53 54 77 | 22 25 26 27 39 |

### 5.6 RSNs definitions in AAL parcellation scheme

### 5.7 Changes in human connectome after modulatory ACh change

**Figure 12:**
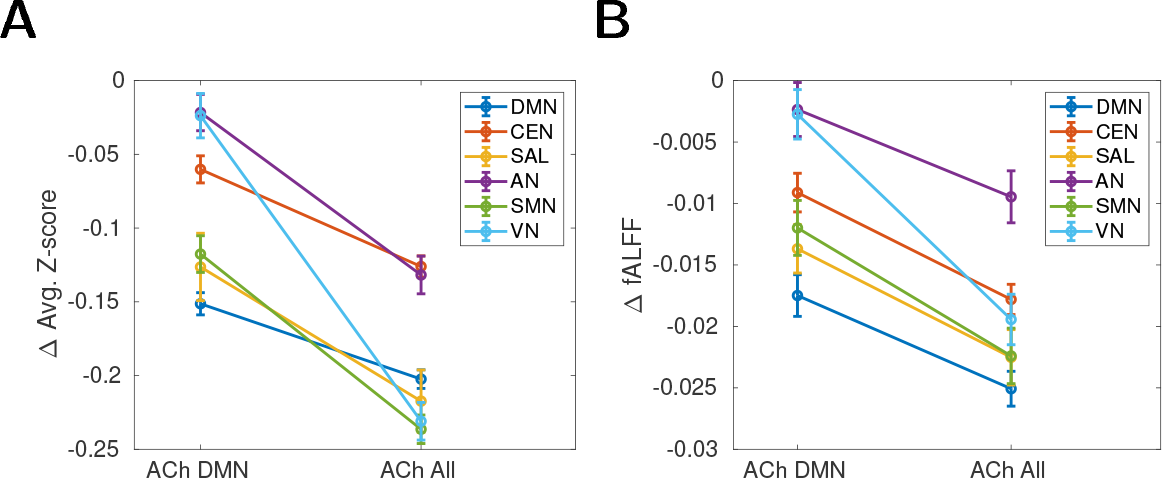
Changes in human connectome after modulatory ACh change. The plots are identical with Fig. 6 A2/A3, except that we removed from all (non-DMN) networks nodes which are part of DMN.

### 5.8 Effect of cholinergic modulation on FC, auxiliary sortings of the AAL nodes

**Figure 13:**
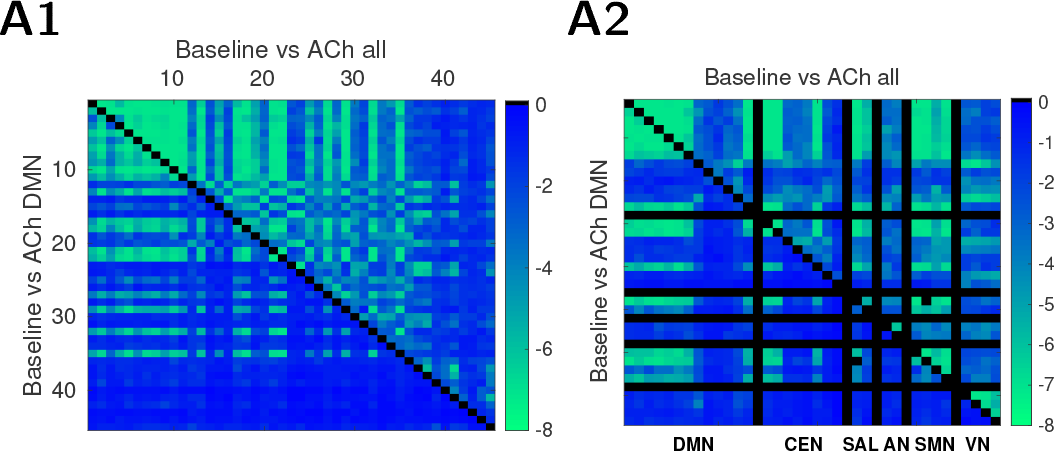
Changes in human connectome after modulatory ACh change. FC changes after ACh release in all (upper triangle) / DMN areas (lower triangle). Color meaning as in Fig. 6 except of sorting of the areas. A1: Areas sorted by their eigenvector-centrality index. A2: Areas regrouped by their affiliation to different resting-state networks, areas in DMN are omitted in other RSNs, so the influence outside of DMN is better visible.

## Notes

### Competing Interest Statement

The authors have declared no competing interest.

### Summary of Updates

Added presentation of results from more detailed model of ACh modulation.

